# Dynamic Causal Modelling Highlights the Importance of Decreased Self-Inhibition of the Sensorimotor Cortex in Motor Fatigability

**DOI:** 10.1101/2023.12.01.569517

**Authors:** Caroline Heimhofer, Marc Bächinger, Rea Lehner, Stefan Frässle, Joshua Henk Balsters, Nicole Wenderoth

**Affiliations:** Neural Control of Movement Lab, Department of Health Sciences and Technology, Zurich Switzerland; Neuroscience Center Zurich (ZNZ), University of Zurich, Federal Institute of Technology Zurich, University and Balgrist Hospital Zurich, Zurich, Switzerland; Translational Neuromodeling Unit, University of Zurich, Swiss Federal Institute of Technology Zurich, Zurich, Switzerland; Department of Psychology, Royal Holloway University of London, Egham, Surrey, United Kingdom; Future Health Technologies, Singapore-ETH Centre, Campus for Research Excellence and Technological Enterprise (CREATE)

**Keywords:** Fatigue, motor slowing, repetitive movements, functional magnetic resonance imaging, dynamic causal modelling

## Abstract

Motor fatigability emerges when challenging motor tasks must be maintained over an extended period of time. It is a frequently observed phenomenon in everyday life which affects patients as well as healthy individuals. Motor fatigability can be measured using simple tasks like finger tapping at maximum speed for 30s. This typically results in a rapid decrease of tapping frequency, a phenomenon called motor slowing. In a previous study (Bächinger et al. 2019), we showed that motor slowing goes hand in hand with a gradual increase of activation in the primary sensorimotor cortex (SM1), supplementary motor area (SMA), and dorsal premotor cortex (PMd). Previous electrophysiological measurements further suggested that the increase in SM1 activity might reflect a breakdown of inhibition and, particularly, a breakdown of surround inhibition which might have led to heightened coactivation of antagonistic muscles. It is unclear what drives the activity increase in SM1 caused by motor slowing and whether motor fatigability affects the dynamic interactions between SM1 and upstream motor areas like SMA and PMd. Here, we performed dynamic causal modelling to answer this question. Our main findings revealed that motor slowing was associated with a significant reduction in SM1 self-inhibition which is in line with previous electrophysiological results. Additionally, the model revealed a significant decrease in the driving input to premotor areas suggesting that structures other than cortical motor areas might cause motor fatigability.

## Introduction

Motor fatigue is a condition which is frequently experienced in everyday life. However, elevated fatigue is a key symptom of many neurological and neuropsychiatric disorders (Kluger, Krupp, and Enoka 2013; Manjaly et al. 2019). Despite its high prevalence in clinical and non-clinical settings, the mechanisms causing motor fatigue are still poorly understood. Here, we focus on one specific aspect of fatigue, called “performance fatigability”, which is defined as an objectively measurable decline in outcome parameters (Kluger, Krupp, and Enoka 2013). Performance fatigability can be studied using a simple finger-tapping paradigm: tapping as quickly as possible for 30s is characterised by a significant decrease in tapping speed, which we refer to as motor slowing (Bächinger et al. 2019). In this paradigm, the tapping speed reflects the objective outcome parameter and the reduction of tapping speed over time is a marker of fatigability. Motor slowing is an interesting paradigm to investigate which brain mechanisms underpin fatigability because neuromuscular and spinal factors have been shown to play only a minor role in mediating the decline of tapping speed over time (Arias et al. 2015; Madrid et al. 2016; 2018; Madinabeitia-Mancebo et al. 2021).

In a recent study, we could show that motor slowing is associated with changes in the sensorimotor network and in the primary sensorimotor cortex in particular (SM1, Bächinger et al. 2019). Despite a decrease in tapping speed, blood-oxygen level dependent (BOLD) activation increased significantly over time in the primary sensorimotor cortex (SM1), dorsal premotor cortex (PMd) and the supplementary motor area (SMA). Neurophysiological measurements revealed that motor slowing is linked to a decrease in inhibition, and specifically surround inhibition within SM1, which correlated with an increase in coactivation of agonistic-antagonistic muscle groups (Bächinger et al. 2019). These findings suggest that motor slowing is associated with the breakdown of inhibitory mechanisms in the primary motor cortex. This proposition is consistent with a population coding model of the primary motor cortex whereby local inhibition can shape the broadness of population tuning curves (Georgopoulos, Schwartz, and Kettner 1986; Georgopoulos and Carpenter 2015): when surround inhibition is low, agonistic and antagonistic tuning curves are wider, resulting in a higher level of coactivation of involved and non-involved muscle groups.

It has been suggested that these local inhibitory mechanisms can be modulated by excitatory connections from other brain areas (Mahan and Georgopoulos 2013). In our previous motor slowing study, we proposed that projections from areas upstream of SM1 may have altered the observed change in inhibition in SM1 (Bächinger et al. 2019). Indeed, our functional magnetic resonance imaging (fMRI) results showed that motor slowing was also characterized by a significant activation increase in dorsal premotor cortex (PMd) and supplementary motor area (SMA). Both areas have been shown to be involved in modulating descending motor commands (see review Correia et al. 2022). It remains unclear, however, how PMd and SMA interact with SM1 while motor slowing arises, and how these areas contribute to shaping local inhibition in SM1.

To investigate premotor-motor interactions during motor slowing, we performed dynamic causal modelling (DCM) on the data from our previous study (Bächinger et al. 2019). Previously, we investigated which brain areas show a change in BOLD activation, but we did not analyse how these brain areas interact. By estimating effective connectivity through patterns of causal interaction, DCM can reveal directionality of neural interactions.

We hypothesized that self-inhibition of SM1 changes during motor slowing which is in line with previous findings that local inhibition is altered with motor slowing. We further investigated whether motor slowing is associated with a change in effective connectivity between SM1, SMA, and PMd. Finally, we explored whether significant changes in effective connectivity correlate with the observed behavioural decrease in movement speed.

## Methods

The data used in this manuscript has been published previously in Bächinger et al. 2019, experiment 6. The focus of the analysis in the previous publication was on changes in fMRI BOLD activation associated with motor slowing. Here, we re-analysed the data with an emphasis on network modelling with DCM. The behavioural task, fMRI preprocessing methods, and the general linear model (GLM) analysis of BOLD activity (i.e., parametric and block design-based analysis) is identical to Bächinger et al. 2019. Relevant parts of the preprocessing are reiterated below for the reader’s convenience.

### Participants

25 participants took part in the experiment and 24 (12 female, mean age: 23.8 +/- 3.3, right-handed) were included in the DCM analysis. One participant was excluded because they did not show any motor slowing, but rather an increase in tapping speed, indicating that they did not perform the task as instructed. All participants were free of medication, had no history of neurological or psychiatric disease and were naïve to the purpose of the experiment. All experimental protocols were approved by the research ethics committee of the canton of Zurich (KEK-ZH 2015-0537) and participants gave written informed consent to the study.

### Behavioural task and analysis

The experiment consisted of two different conditions: finger tapping for either 30 s (slowing condition) or 10 s (control condition), each followed by a 30 s break. Tapping was performed alternating between index and middle finger at maximum speed. Participants were informed about the condition prior to the start of tapping with a visual get-ready cue (randomly jittered between 2-3 s). The conditions were blocked within each fMRI run: One block was made up of four trials of the slowing condition, followed by four trials of the control condition, or vice versa. Each participant performed two fMRI runs consisting of two blocks each. This resulted in 16 trials per condition. The starting condition of the first run (slowing or control condition) was alternated across participants and the second run had a counterbalanced order in relation to the first run. Additionally, an implicit baseline of 20 s was measured after each block (Figure 1A). Behavioural data was analysed as described previously (Bächinger et al. 2019). Tapping and break intervals were divided into 10 s bins and movement speed was normalised to the average speed of the control condition per participant. This normalised movement speed was subjected to a linear mixed effects model with the fixed factor *time* (i.e., time bins) and the random factor *participant*. Motor slowing was defined as a significant main effect of *time* (Figure 1B).

**Figure 1:**
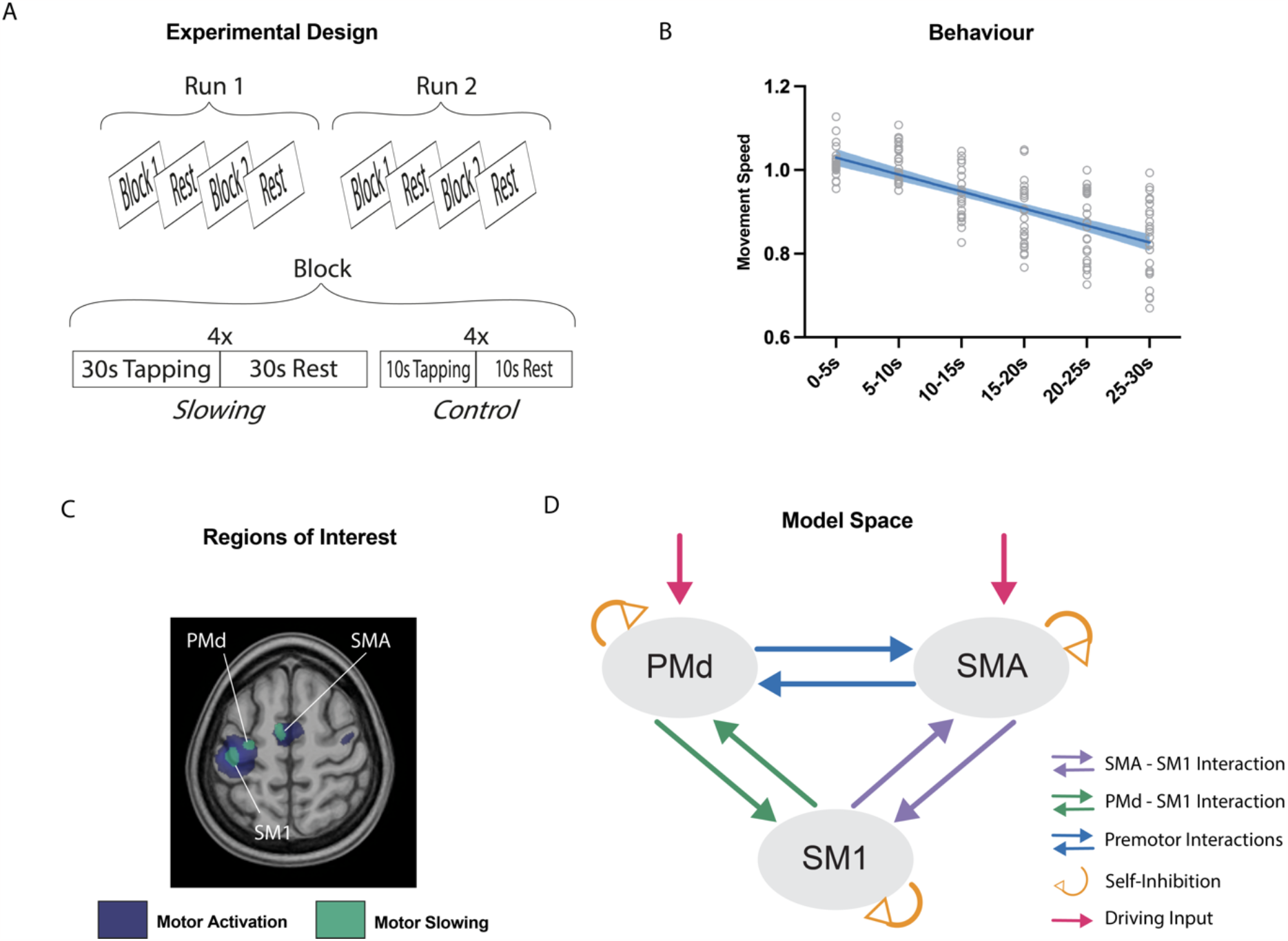
(A) Experimental Design of fMRI study. 24 participants were either tapping for 30s (slowing condition) or 10s (control condition) during fMRI scanning. (B) Behavioural Results. Behavioural results showing a significant decrease in movement speed over 30s of tapping. (C) fMRI activations, associated with either tapping itself (motor network, blue) or motor slowing (increasing activation with decreasing movement speed, green). Regions of interest where defined based on the closest individual activations of PMd, SMA, and SM1. (D) Schematic overview of the model space for DCM.

### fMRI acquisition and preprocessing

fMRI scans were acquired with a Philips Ingenia 3T whole body scanner. Prior to the functional runs, high resolution T1-weighted anatomical scans were acquired (voxel size = 1 mm^3^, 160 sagittal slices, matrix size = 240x240, TR/TE = 8.3/3.9 ms). These anatomical scans were used for functional image registration and normalisation. During the behavioural runs 360 volumes were acquired in each run (voxel size = 2.75x2.75x3.3 mm, matrix size = 128x128, TR/TE = 2500/35 ms, flip angle = 82 degrees, 40 slices acquired in interleaved order for full brain coverage). Preprocessing was performed using SPM12 (Wellcome Trust) with default parameters and consisted of realignment to the average functional image, segmentation of the anatomical image, normalization to MNI space, and spatial smoothing (8 mm isotropic Gaussian kernel at full-width-half maximum).

### fMRI data analysis

fMRI analyses were also performed in SPM12. The first-level model of each participant consisted of a general linear model. The GLM design matrix included four regressors of interest: tapping, parametric modulation of tapping, recovery, and parametric modulation of recovery. The tapping regressor represented the time periods when the participant was tapping. The recovery reflects the 30 s rest condition after a tapping trial. The parametric modulation regressor consisted of a linear increase over the tapping periods (reflecting the increase in motor slowing) or a linear increase over the recovery period after a 30 s tapping trial (but not a 10 s tapping trial). The linear increase was the same across all participants and did not depend on the participant’s performance. Importantly, the parametric modulation regressor was orthogonalized with respect to the tapping regressor. Note that the 30 s slowing condition and the 10 s control condition were modelled together in each regressor. For the parametric modulator, the slowing condition consisted of a linear increase in six bins of 5 s, and the control condition was made up of a linear increase in two bins of 5 s. Regressors of no interest in the GLM consisted of get-ready periods and six head movement parameters (translation and rotation along the x, y, and z-axis). All regressors except the six head movement parameters were convolved with a canonical hemodynamic response function. The two regressors of interest were contrasted against the implicit baseline and were then subjected to a second-level random-effects analysis across participants. The second level analysis was a single one-sample t-test contrasting the regressors of interest against zero. P-values smaller than 0.05 family-wise error (FWE) corrected for multiple comparisons were considered statistically significant. Localisation of functional clusters was aided by the anatomy toolbox (Eickhoff et al. 2005).

### Dynamic causal modelling

To investigate the changes in effective (directed) connectivity with motor slowing, we performed DCM (Friston, Harrison, and Penny 2003) using SPM12. In short, dynamic causal models are generative models that aim to capture directed interactions among brain regions or states based on à priori hypotheses. In DCM, changes in brain states over time are modelled in the form of a state-space equation:

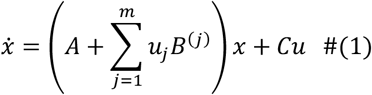

In this equation, *x* is the state vector representing the current neuronal state and 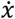 refers to the change in the neuronal state over time. The matrix *A* represents the underlying endogenous or intrinsic connectivity with fixed weights defined by the model, whereas *B*^*(j)*^ reflects the weights of task-dependent modulations of connectivity, driven by external modulatory inputs *u*_*j*_. *C* represents the weights for direct inputs, characterising how the extrinsic driving input *u* directly influence brain regions.

In the scope of this study, the matrix *A* represents the endogenous connectivity of the motor system (i.e., bidirectional connectivity between SM1, PMd, and SMA; see also regions of interests below). The term *u*_*j*_*B*^(*j*)^characterises the strength of the modulatory changes that occur due to motor slowing, which are modelled as a linear increase reflecting motor slowing. The final term *Cu*, describes the external driving input to the (pre-) motor system, modelled here as a constant input from prefrontal areas to either PMd, SMA, or both.

### Regions of interest, endogenous connectivity, and its modulation

As a hypothesis driven method, DCM requires a neurobiologically-plausible model of connectivity to be defined à priori. We therefore selected regions of interest which were associated with motor slowing. Specifically, we previously found that activity (pFWE < 0.05) of the left SM1, left PMd, and bilateral SMA (Bächinger et al. 2019) were inversely correlated with motor slowing of the right hand: All these regions showed an activation increase with decreasing tapping speed. Based on this finding, we investigated here whether motor slowing is associated with changes in premotor-motor interactions. To that end we built several DCMs incorporating PMC, SMA, and SM1 (i.e., the three areas directly associated with motor slowing).

We extracted the BOLD signal time-series of our Effect of Interest from 4mm radius spheres centred on the following three regions of interest: SM1, PMd, and SMA. The Effect of Interest consisted of our four regressors of interest (tapping, parametric modulation of tapping, recovery, parametric modulation of recovery). All regions of interest were defined by taking the coordinates from the group-level analysis (Supplementary Material 1) and then extracting the closest peak-level activation on the single subject level.

The endogenous connectivity matrix (matrix *A* in Equation (1)) was defined by previous anatomical studies: Specifically, we assumed that all regions are connected bidirectionally based on previous anatomical findings (Luppino et al. 1993; Rouiller et al. 1994; as cited in Michely et al. 2015). Also, all included regions were assumed to be self-modulatory. Self-modulations were chosen to represent two-state models and they can therefore be considered to be self-inhibitory (Marreiros, Kiebel, and Friston 2008).

The extrinsic regressor (term *u* in Equation (1)) that modulates connectivity of the network (term *B* in Equation (1)) reflected the effect of motor slowing. In the scope of this study, motor slowing was simplified as a linear change over time, as represented by the parametric modulation (see section *fMRI data analysis*) which served as input for this analysis. The driving input to the model was assumed to be a constant input from prefrontal areas (Michely et al. 2015).

### Model space and model families

As our main interest was to test whether self-inhibition of SM1 is crucial to explaining modulation of connectivity during motor slowing, we split the model space into model families with and without self-inhibition of SM1. Further, we wanted to investigate whether the premotor areas (SMA, PMd or both) shape the self-inhibition in SM1. Therefore, we set up multiple model families: (1) The top-down model family (in accordance with our main hypothesis outlined in the introduction), in which connections from SMA to SM1 and from PMd to SM1 were modulated. (2) The bottom-up model family, to verify whether our hypothesis may be inversed, meaning self-inhibition of SM1 may modulate SMA and PMd in a bottom-up fashion. In this model family, connections from SM1 to SMA and from SM1 to PMd were modulated. (3) The selective premotor model family, to test whether one premotor area is much more strongly involved in modulations of motor slowing: only SM1-SMA or SM1-PMd connections were modulated in these models. (4) The null model family. In this model family, none of the connections between any of the premotor areas and SM1 were modulated.

As mentioned, our main interest was to determine the necessity of self-inhibition in SM1, which is why these 4 model families were further specified as either having self-inhibition of SM1 modulated or not. This resulted in 8 model families: Top-down models without self-inhibition of SM1 (36 models, Figure 2A), bottom-up models without self-inhibition of SM1 (36 models, Figure 2B), selective premotor models without self-inhibition of SM1 (12 models, Figure 2C), null models without self-inhibition of SM1 (6 models, Figure 2D), top-down models with self-inhibition of SM1 (36 models, Figure 2E), bottom-up models with self-inhibition of SM1 (36 models, Figure 2F), selective premotor models with self-inhibition of SM1 (12 models, Figure 2G), null models with self-inhibition of SM1 (6 models, Figure 2H).

**Figure 2:**
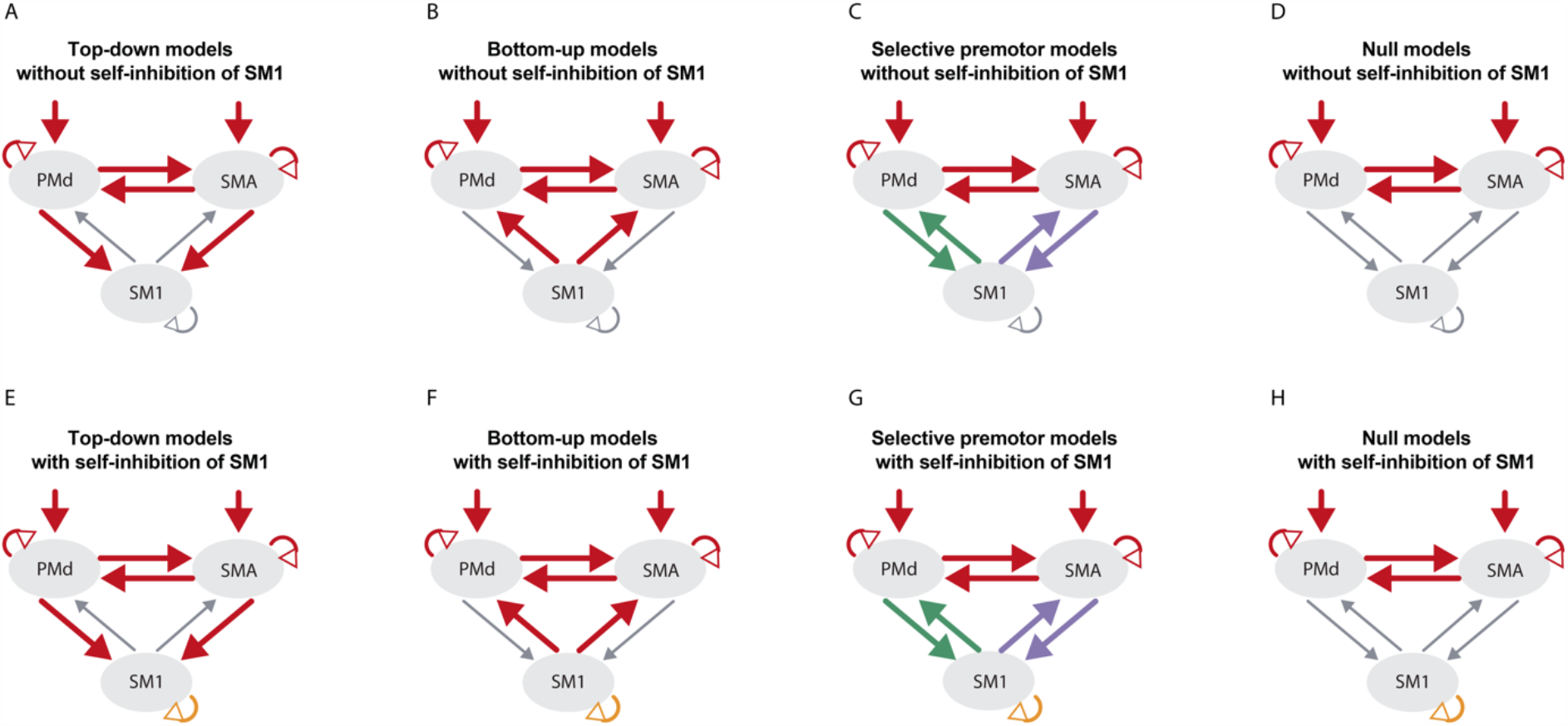
Model Families. One model family consisted of several models differing in which connections were modulated. The connections that may have been modulated within one model family are marked in red. The selective premotor models (C and G) either had modulations of the SMA-SM1 (purple) or the PMd-SM1 (green) interactions, but not both in the same model. The premotor interaction (SMA to PMd and PMd to SMA) were always modelled together. The driving input was set either to PMd, SMA, or both. Self-inhibition of SMA and PMd were only modulated in combination with a modulated connection with SM1. However, for model families A and E, the premotor-motor connections were also modulated without self-inhibition of the corresponding premotor area. Self-inhibition of SM1 (orange) was assumed in all models of model families E-H.

In all model families, the models were set up (i) with and without modulation of premotor interactions between SMA and PMd (Figure 1D, blue), and (ii) with the driving input set to either PMd, SMA, or both (Figure 1D, red). The main modulations of the selective premotor model family were either a bidirectional modulation of SM1-SMA (Figure 1D, purple) or SM1-PMd (Figure 1D, green). In both cases, the involved premotor area was modelled with self-inhibition (Figure 1D, orange). The top-down and bottom-up model family differed in whether the connections SM1-PMd, SM1-SMA, or both were modulated, with or without self-inhibition of the involved premotor area. All in all, the model space consisted of 180 models, split into 8 model families. A list of all the models can be found in Supplementary Material 2.

#### Model selection and statistical analysis

To identify the most likely model family given the data, we used random-effects family-level Bayesian model selection (Penny et al. 2004; Stephan et al. 2009). As the model selection did not reveal decisive evidence for a single winning model family (family exceedance probabilities < 0.95, Figure 3), all models across all model families were averaged through Bayesian model averaging (BMA) with an Occam’s window of 0.05 to inspect the model parameters. These BMA parameter estimates were then subjected to two further analyses (Stephan et al. 2010). First, to identify the connections which were significantly modulated by motor slowing across participants, a group level post-hoc analysis on the maximum-à-posteriori (MAP) of the matrix *B* was performed using Bonferroni-corrected t-tests. Secondly, a stepwise linear regression was performed to identify which of these modulated connections were directly associated with individual differences in motor slowing as quantified by the behavioural data. The regression model tested whether behavioural changes in tapping speed can be explained by the MAPs of the modulated connections.

**Figure 3:**
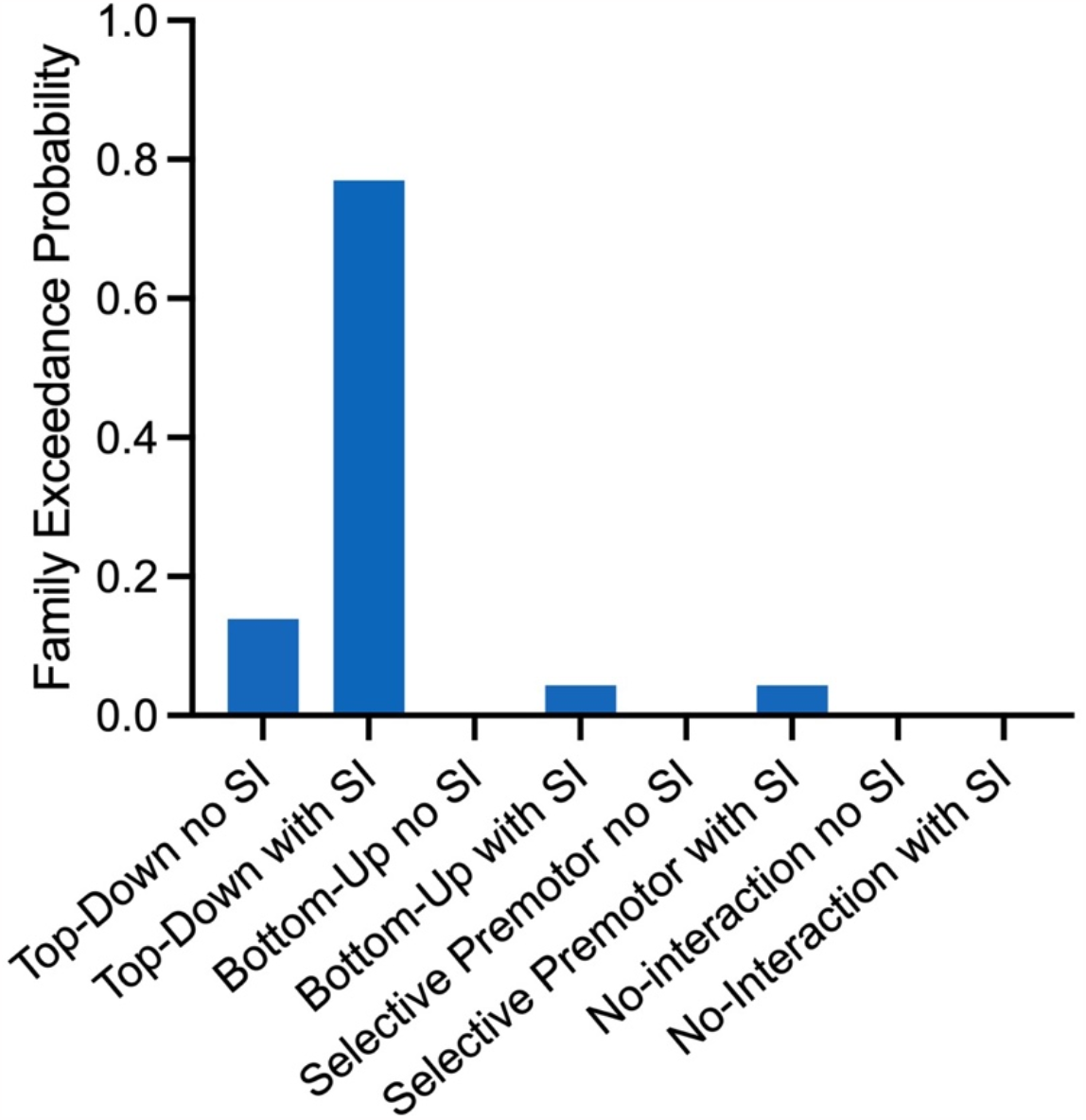
Exceedance probabilities of random-effects Bayesian model selection for defining winning model family. No model family reached the evidence threshold (> 0.95).

## Results

With DCM, we investigated if motor slowing is associated with changes in effective connectivity in the motor network. We first ran family-wise random-effects Bayesian model selection (Penny et al. 2010), which revealed the highest probability for the top-down model family with self-inhibition of SM1. The variance explained by the models of this model family is on average 31.2% (see Supplementary Material 3 for variance explained per participant). However, evidence for this model family was not decisive (i.e., exceedance probability < 0.95) (Figure 3). Therefore, we performed Bayesian model averaging across the whole model space. The variance explained by models in the Occam’s window, which are the ones that were averaged, can be found in Supplementary Material 4. The BMA revealed that motor slowing was accompanied by decreased connectivity in driving inputs to PMd and SMA, decreased connectivity from PMd to SM1, and decreased self-inhibition of SMA, as well as SM1. An increase in connectivity was found bidirectionally between PMd and SMA, bidirectionally between SMA and SM1, unidirectionally from SM1 to PMd, and for self-inhibition of PMd (Figure 4).

**Figure 4:**
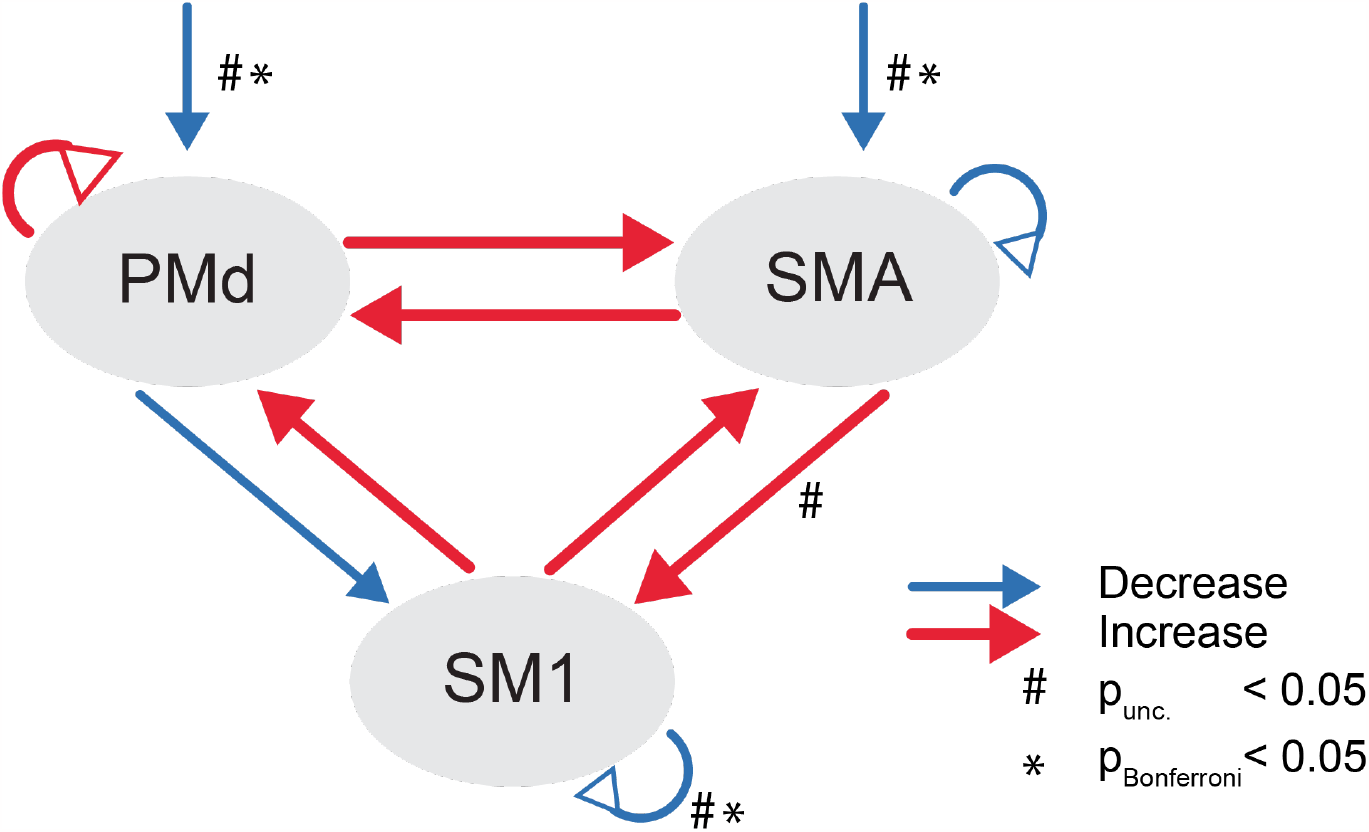
BMA model results. Arrows with triangle heads represent (self-)inhibitory connectivity, the other arrows represent facilitatory connectivity. Increase in connectivity or self-inhibition with increasing motor slowing are shown in red, decrease in connectivity or self-inhibition with increasing motor slowing are shown in blue. The arrows towards PMd and SMA reflect the constant driving input over the whole tapping period. #<0.05 uncorrected, *p<0.05 Bonferroni corrected

To identify which of these connections were significantly modulated during motor slowing, we tested the maximum-à-posteriori (MAPs) estimates against 0 (one-sample t-test with Bonferroni correction). Significant modulation was only found for the decrease of self-inhibition of SM1 and the decrease in driving inputs to the premotor areas (Figure 4, *pBonferroni < 0.05). There was a trend that effective connectivity from SMA to SM1 increased with motor slowing but this effect did not survive Bonferroni correction.

In an additional analysis, we tested whether any changes in effective connectivity were directly associated with the behavioural effects of motor slowing. In a stepwise linear regression analysis, we found that stronger connectivity from SMA to SM1 and from SM1 to SMA were linearly associated with more motor slowing (adjusted R^2^=0.514; p<0.01). However, partial residual analysis revealed that these associations could have been driven by outlier values (Supplementary Material 5).

## Discussion

High motor fatigability, as reflected by the phenomenon of motor slowing, is characterised by increased activity within the cortical sensorimotor network. This observation is somewhat paradoxical, since higher cortical BOLD activity has been associated with higher tapping speed when tested in a non-fatigued state (Rao et al. 1996; Schlaug et al. 1996; Sadato et al. 1997; Jäncke, Specht, et al. 1998; Jäncke, Peters, et al. 1998; Deiber et al. 1999; Agnew, Zeffiro, and Eden 2004; Lutz et al. 2004). Here, we used DCM to test how motor slowing modulates the interaction between PMd, SMA, and SM1. Our main result shows that motor slowing was characterised by a significant decrease of (i) SM1 self-inhibition, and (ii) excitatory driving inputs to both premotor areas (SMA and PMd). We further observed an increase in SMA to SM1 connectivity at a trend level.

DCM revealed that self-inhibition of SM1 decreased during motor slowing. Interestingly, the driving inputs to the premotor areas decreased significantly, while interactions between premotor and motor areas only reached trend level significance for a connectivity increase from SMA to SM1. This suggests that the release of self-inhibition in SM1 is the major contributor to the increase of BOLD activity typically observed during motor slowing. This finding is in line with our previous neurophysiological results (Bächinger et al. 2019), but also with a larger body of research that used different fatiguing paradigms to show that intracortical inhibition is reduced after fatiguing muscle contractions (Benwell et al. 2006; Maruyama et al. 2006; Hunter et al. 2016; Latella et al. 2020).

We further found that the driving input to the cortical premotor areas decreases during motor slowing. Despite this finding being consistent with the reduction in tapping speed, the decrease in facilitatory input makes it unlikely that influences from other areas drive the high activity in SM1, PMd, and SMA. This is interesting because the driving input to premotor areas seems to play a more important role for motor slowing than premotor-primary motor interactions. Where these driving inputs originate has yet to be defined, but basal ganglia, thalamus, or the cerebellum are likely candidates based on previous research (Liu et al. 2003; Van Duinen et al. 2007; Post et al. 2009; Hou et al. 2016; Bächinger et al. 2019). The cerebellum is particularly interesting, as it has been found to be involved in the continuous monitoring of movement rates, as well as in regulating movement rhythmicity via the brain stem, basal ganglia, and thalamus (Scott 2004; Bastian 2006; Pisotta and Molinari 2014; Therrien and Bastian 2019).

Concerning premotor-motor interactions, we found a trend-level increase in effective connectivity from SMA to SM1 with motor slowing, and the stepwise linear regression analysis hinted that stronger effective connectivity between SMA and SM1 was associated with more slowing on an individual level. SMA is known for its importance of controlling internally generated movements through projections to M1 (Samuel 1997; Konoike and Nakamura 2020). In this regard, SMA has not only been linked to the temporal sequencing of movements (Tanji 2001), but also to rhythm production itself (Konoike and Nakamura 2020). Increased SMA activity has been proposed to not be related to the execution of movement per se, but rather to the elevated difficulty in motor control (Kawashima et al. 1999). In other fatiguing paradigms, SMA has been linked to effort perception (Zénon, Sidibé, and Olivier 2014; Sharples et al. 2016; Emanuel et al. 2021), with modulation of SMA leading to changes in perceived effort. This suggests that the increased SMA to SM1 connectivity that we found may reflect the increase in perceived effort required to maintain the tapping performance with advancing time. However, this conclusion is highly speculative and must be interpreted with caution since the association between SMA and behaviour has been driven by a small group of participants and we did not measure the perceived effort. Thus, even though interactions between premotor and primary sensorimotor cortex might contribute to motor slowing, it seems likely that more complex neuronal interactions involving a larger network may underlie the observed behavioural outcome.

In summary, these results revealed additional evidence supporting the hypothesis that motor slowing is associated with a release of inhibition in SM1. Additionally, the driving input coming from other (motor) areas is essential for explaining the network modulations occurring during motor slowing.

## Conclusion

Our DCM analysis indicates that a reduction in self-inhibition of SM1 explains the increase in BOLD activation that occurs during motor slowing, even though it is not directly associated with the behavioural decrease in movement speed. Furthermore, we show that premotor-motor interactions are moderately modulated by motor slowing, but that the driving input to the cortical motor network appears to play an even more important role. These findings emphasize that processes upstream of the premotor-motor areas contribute to the changes occurring in SM1 when fatigability increases during fast finger tapping.

## Acknowledgements

The authors would like to thank Samira Hanimann for data collection and Ingrid Odermatt, Jenny Imhof, and Daniel Woolley for proof-reading the manuscript.

## Author Contributions

Caroline Heimhofer: Data curation, Software, Formal analysis, Investigation, Visualization, Methodology, Writing—original draft, Project administration, Writing—review and editing; Marc Bächinger: Conceptualization, Resources, Data curation, Software, Formal analysis, Investigation, Visualization, Methodology, Writing— original draft, Project administration, Writing—review and editing; Rea Lehner: Conceptualization, Resources, Methodology, Formal analysis, Supervision, Writing—original draft; Stefan Frässle: Software, Methodology, Formal analysis, Supervision, Writing—review and editing; Joshua Henk Balsters: Conceptualization, Formal analysis, Supervision; Nicole Wenderoth: Conceptualization, Formal analysis, Supervision, Funding acquisition, Methodology, Writing—original draft, Project administration, Writing—review and editing

## Supplementary Material

**Supplementary Material 1.**
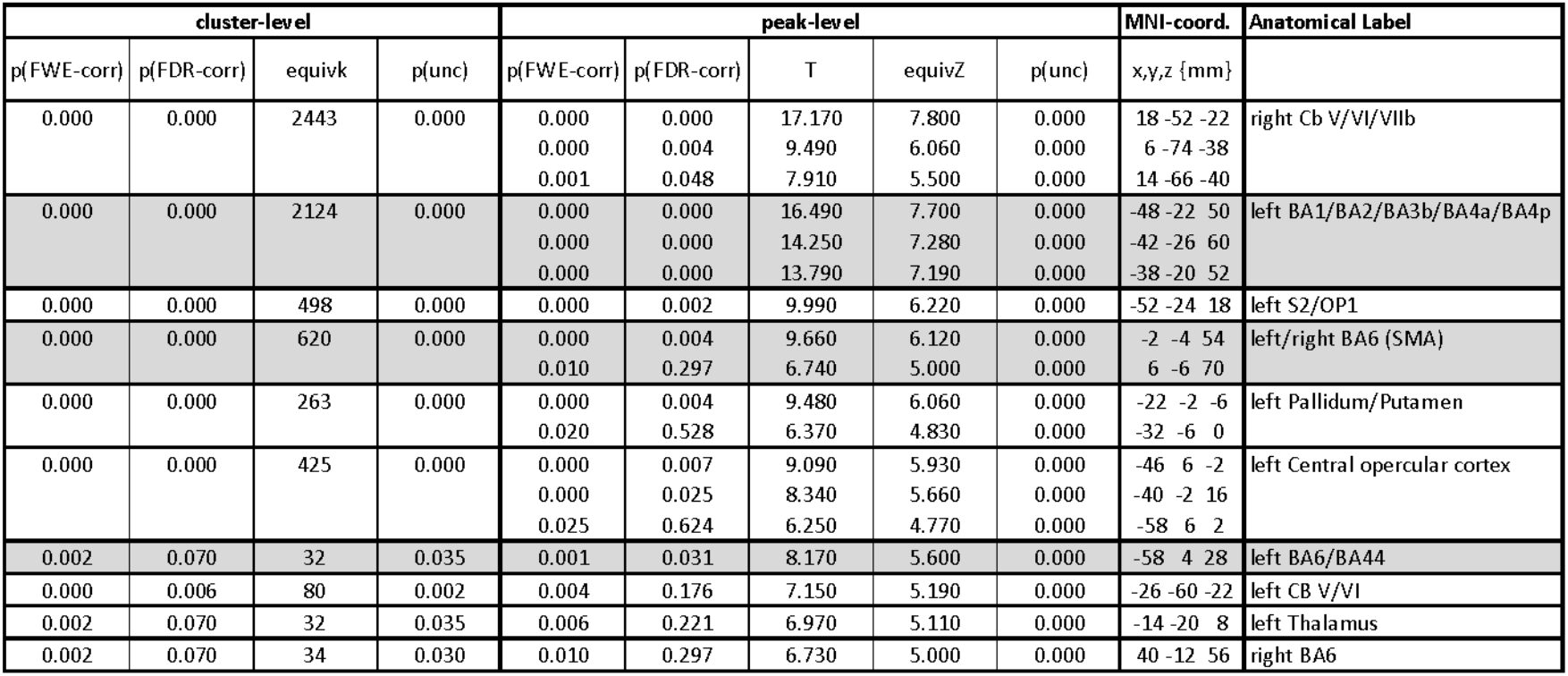
Peak activations during tapping. Areas highlighted in grey were associated with motor slowing and used as regions of interest in the DCM analysis.

| cluster-level |  |  |  | peak-level |  |  |  |  | MNI-coord. | Anatomical Label |
| --- | --- | --- | --- | --- | --- | --- | --- | --- | --- | --- |
| p(FWE-corr) | p(FDR-corr) | equivk | p(unc) | p(FWE-corr) | p(FDR-corr) | T | equivZ | p(unc) | x,y,z {mm} |  |
| 0.000 | 0.000 | 2443 | 0.000 | 0.000 | 0.000 | 17.170 | 7.800 | 0.000 | 18 -52 -22 | right Cb V/VI/VIIb |
|  |  |  |  | 0.000 | 0.004 | 9.490 | 6.060 | 0.000 | 6 -74 -38 |  |
|  |  |  |  | 0.001 | 0.048 | 7.910 | 5.500 | 0.000 | 14 -66 -40 |  |
| 0.000 | 0.000 | 2124 | 0.000 | 0.000 | 0.000 | 16.490 | 7.700 | 0.000 | -48 -22 50 | left BA1/BA2/BA3b/BA4a/BA4p |
|  |  |  |  | 0.000 | 0.000 | 14.250 | 7.280 | 0.000 | -42 -26 60 |  |
|  |  |  |  | 0.000 | 0.000 | 13.790 | 7.190 | 0.000 | -38 -20 52 |  |
| 0.000 | 0.000 | 498 | 0.000 | 0.000 | 0.002 | 9.990 | 6.220 | 0.000 | -52 -24 18 | left S2/OP1 |
| 0.000 | 0.000 | 620 | 0.000 | 0.000 | 0.004 | 9.660 | 6.120 | 0.000 | -2 -4 54 | left/right BA6 (SMA) |
|  |  |  |  | 0.010 | 0.297 | 6.740 | 5.000 | 0.000 | 6 -6 70 |  |
| 0.000 | 0.000 | 263 | 0.000 | 0.000 | 0.004 | 9.480 | 6.060 | 0.000 | -22 -2 -6 | left Pallidum/Putamen |
|  |  |  |  | 0.020 | 0.528 | 6.370 | 4.830 | 0.000 | -32 -6 0 |  |
| 0.000 | 0.000 | 425 | 0.000 | 0.000 | 0.007 | 9.090 | 5.930 | 0.000 | -46 6 -2 | left Central opercular cortex |
|  |  |  |  | 0.000 | 0.025 | 8.340 | 5.660 | 0.000 | -40 -2 16 |  |
|  |  |  |  | 0.025 | 0.624 | 6.250 | 4.770 | 0.000 | -58 6 2 |  |
| 0.002 | 0.070 | 32 | 0.035 | 0.001 | 0.031 | 8.170 | 5.600 | 0.000 | -58 4 28 | left BA6/BA44 |
| 0.000 | 0.006 | 80 | 0.002 | 0.004 | 0.176 | 7.150 | 5.190 | 0.000 | -26 -60 -22 | left CB V/VI |
| 0.002 | 0.070 | 32 | 0.035 | 0.006 | 0.221 | 6.970 | 5.110 | 0.000 | -14 -20 8 | left Thalamus |
| 0.002 | 0.070 | 34 | 0.030 | 0.010 | 0.297 | 6.730 | 5.000 | 0.000 | 40 -12 56 | right BA6 |

**Supplementary Material 2.**
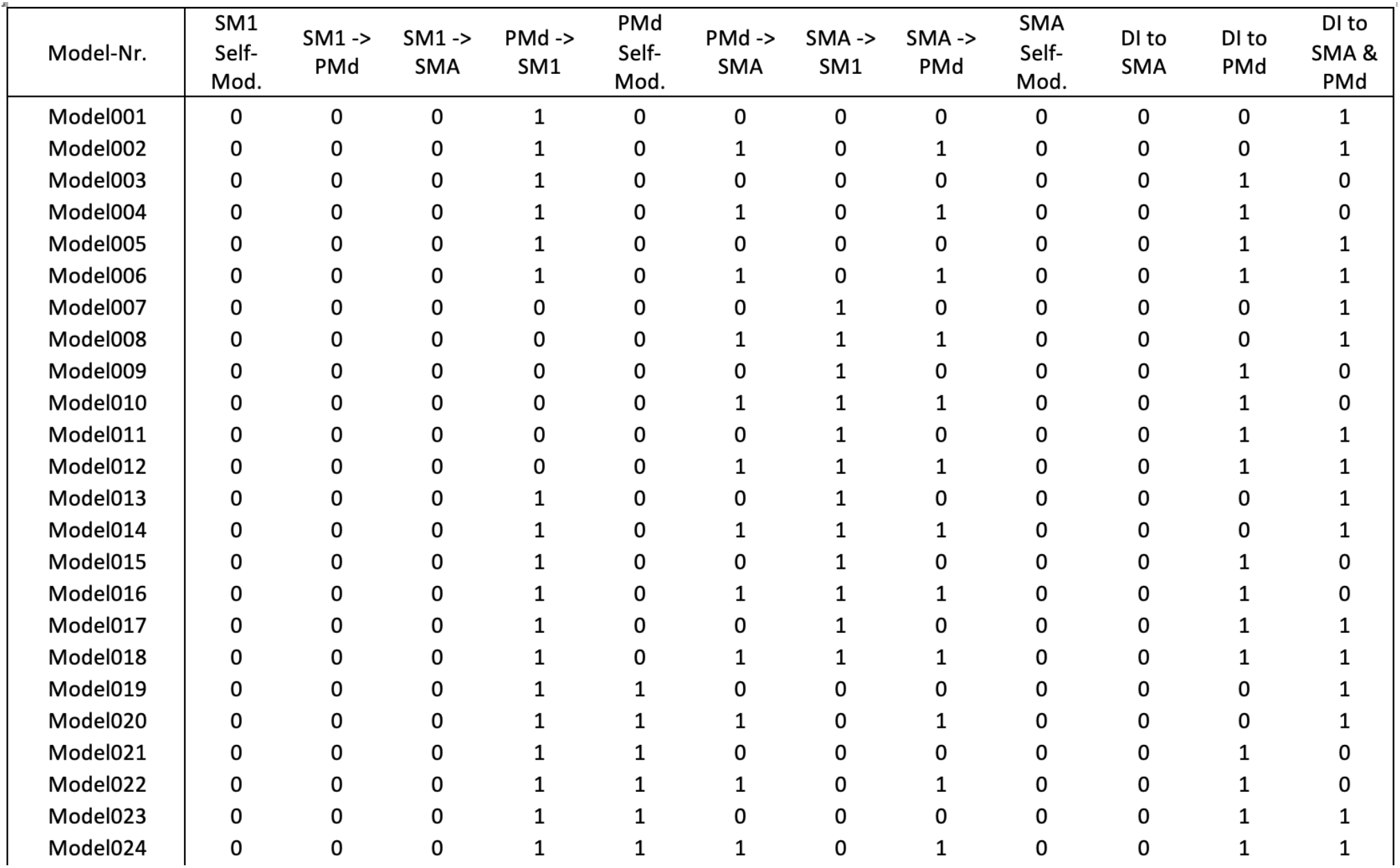
*Model space. The first six columns indicate whether motor slowing had a modulatory effect on the respective connections. The last three columns reflect to which nodes (SMA, PMd, or both) the driving input (DI) of tapping was set. Corresponding model families are listed here: Models 1-36: top-down models without self-inhibition of SM1; Models 37-72: top-down models with self-inhibition of SM1; Models 73-108: bottom-up models without self-inhibition of SM1; Models 109-144: bottom-up models with self-inhibition of SM1; Models 145-156:* selective premotor *models without self-inhibition of SM1; Models 157-168:* selective premotor *models with self-inhibition of SM1; Models 196-174: null models without self-inhibition of SM1; Models 175-180: null models with self-inhibition of SM1*.

**Supplementary Material 3.**
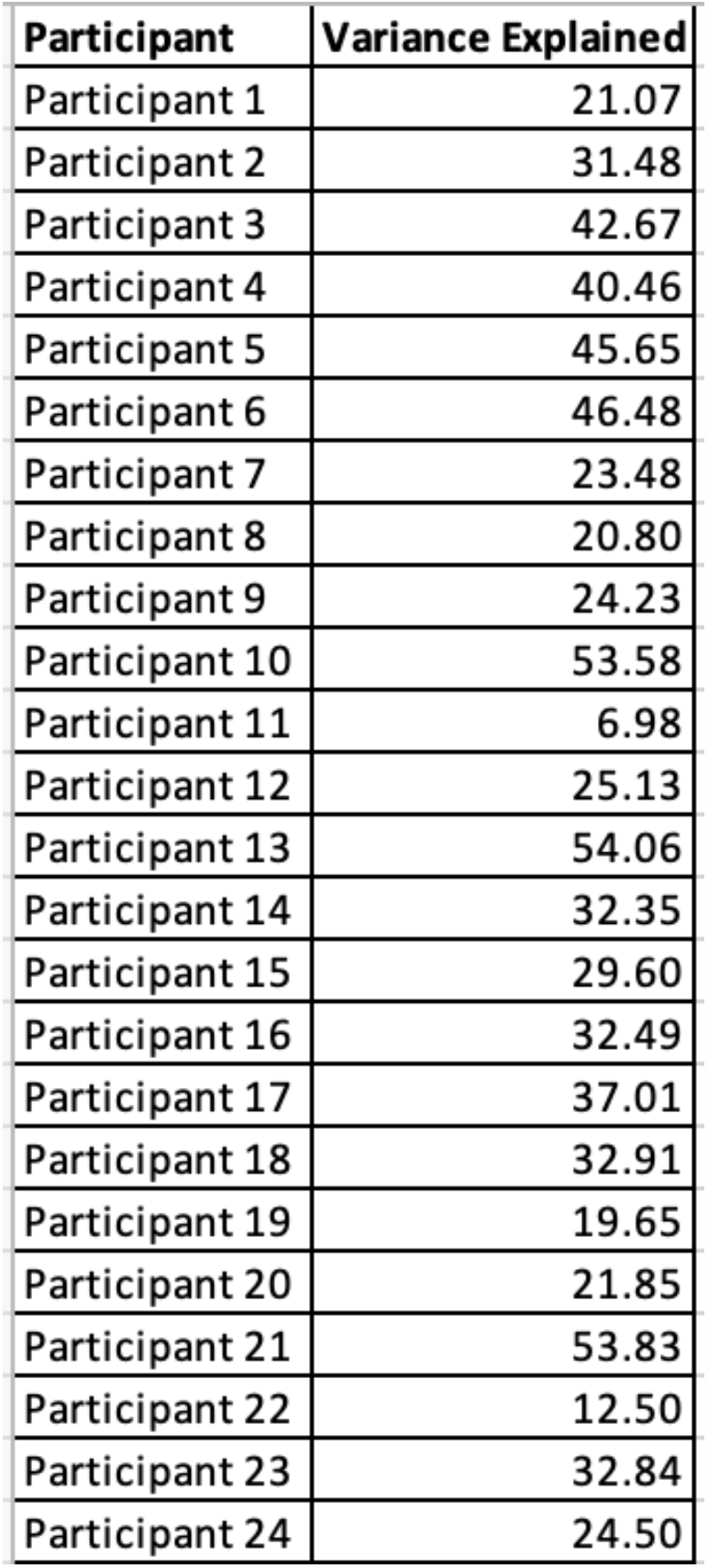
Variances explained by models of winning model family per participant. The variances explained by the models of the winning model family were averaged per participant.

| Participant | Variance Explained |
| --- | --- |
| Participant 1 | 21.07 |
| Participant 2 | 31.48 |
| Participant 3 | 42.67 |
| Participant 4 | 40.46 |
| Participant 5 | 45.65 |
| Participant 6 | 46.48 |
| Participant 7 | 23.48 |
| Participant 8 | 20.80 |
| Participant 9 | 24.23 |
| Participant 10 | 53.58 |
| Participant 11 | 6.98 |
| Participant 12 | 25.13 |
| Participant 13 | 54.06 |
| Participant 14 | 32.35 |
| Participant 15 | 29.60 |
| Participant 16 | 32.49 |
| Participant 17 | 37.01 |
| Participant 18 | 32.91 |
| Participant 19 | 19.65 |
| Participant 20 | 21.85 |
| Participant 21 | 53.83 |
| Participant 22 | 12.50 |
| Participant 23 | 32.84 |
| Participant 24 | 24.50 |

**Supplementary Material 4.** Variances explained by models in the Occam’s window per participant. The model number with the variance explained for this participant are listed. The number of models in the Occam’s window varied across participant.

| Participant | model_1 | model_2 | model_3 | model_4 | variance explained_1 | variance explained_2 | variance explained_3 | variance explained_4 |
| --- | --- | --- | --- | --- | --- | --- | --- | --- |
| Participant 1 | 50 | 68 |  |  | 27.23 | 28.27 |  |  |
| Participant 2 | 126 |  |  |  | 34.59 |  |  |  |
| Participant 3 | 174 |  |  |  | 43.74 |  |  |  |
| Participant 4 | 34 |  |  |  | 43.7 |  |  |  |
| Participant 5 | 70 | 72 |  |  | 46.01 | 46.25 |  |  |
| Participant 6 | 106 | 124 | 142 |  | 46.62 | 46.7 | 46.76 |  |
| Participant 7 | 34 | 69 | 70 | 106 | 24.44 | 24.57 | 24.79 | 24.59 |
| Participant 8 | 52 |  |  |  | 39.5 |  |  |  |
| Participant 9 | 158 |  |  |  | 39.44 |  |  |  |
| Participant 10 | 16 |  |  |  | 56.97 |  |  |  |
| Participant 11 | 16 |  |  |  | 23.5 |  |  |  |
| Participant 12 | 144 |  |  |  | 31.41 |  |  |  |
| Participant 13 | 164 |  |  |  | 57.78 |  |  |  |
| Participant 14 | 36 | 52 | 72 |  | 34.73 | 34.54 | 34.87 |  |
| Participant 15 | 166 |  |  |  | 40.2 |  |  |  |
| Participant 16 | 16 | 52 | 69 |  | 36.94 | 36.79 | 36.85 |  |
| Participant 17 | 52 | 60 | 160 |  | 43.71 | 43.18 | 41.38 |  |
| Participant 18 | 166 |  |  |  | 32.88 |  |  |  |
| Participant 19 | 126 |  |  |  | 28.91 |  |  |  |
| Participant 20 | 72 |  |  |  | 24.23 |  |  |  |
| Participant 21 | 60 | 72 |  |  | 54.78 | 54.76 |  |  |
| Participant 22 | 68 |  |  |  | 12.53 |  |  |  |
| Participant 23 | 16 | 52 | 70 |  | 33.83 | 33.8 | 34.03 |  |
| Participant 24 | 34 | 70 | 106 | 142 | 30.63 | 29.95 | 29.46 | 30.36 |

**Supplementary Material 5.**
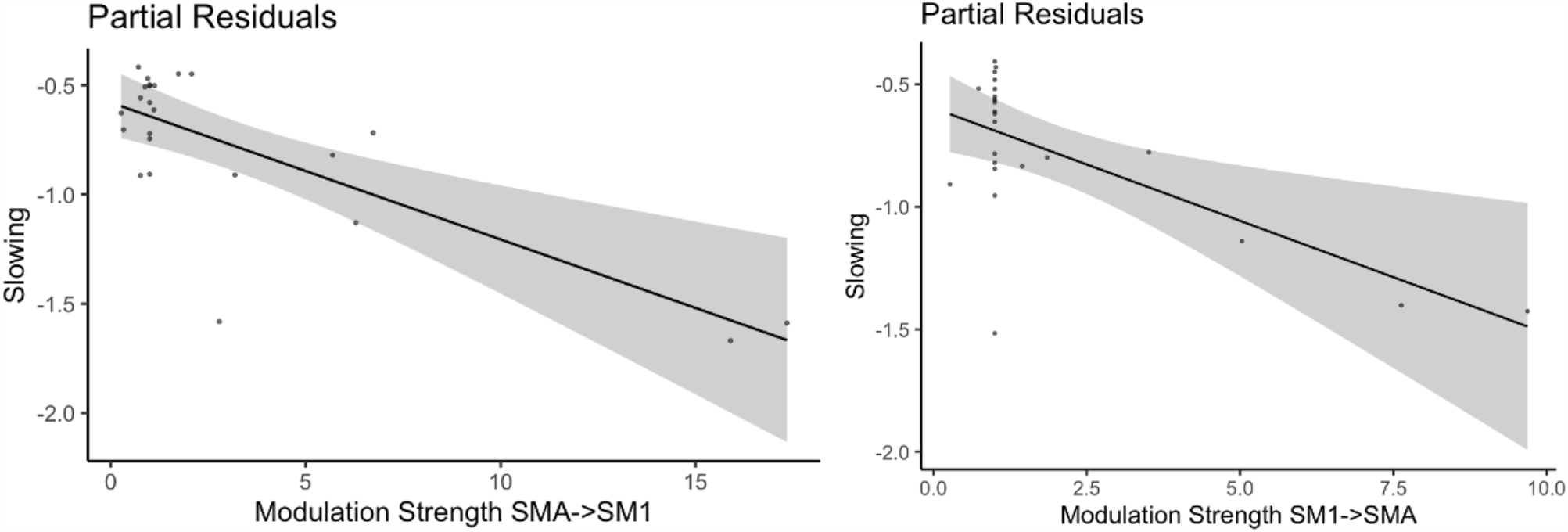
Partial residuals of stepwise linear regression analysis. Both significant regressors seem to be driven by outliers.

## References

Agnew, John A, Thomas A Zeffiro, and Guinevere F Eden. 2004. ‘Left Hemisphere Specialization for the Control of Voluntary Movement Rate’. NeuroImage 22 (1): 289–303. 10.1016/j.neuroimage.2003.12.038.

Arias, P., V. Robles-García, Y. Corral-Bergantiños, A. Madrid, N. Espinosa, J. Valls-Solé, L. Grieve, A. Oliviero, and J. Cudeiro. 2015. ‘Central Fatigue Induced by Short-Lasting Finger Tapping and Isometric Tasks: A Study of Silent Periods Evoked at Spinal and Supraspinal Levels’. Neuroscience 305 (October): 316–27. 10.1016/j.neuroscience.2015.07.081.

Bächinger, Marc, Rea Lehner, Felix Thomas, Samira Hanimann, Joshua Balsters, and Nicole Wenderoth. 2019. ‘Human Motor Fatigability as Evoked by Repetitive Movements Results from a Gradual Breakdown of Surround Inhibition’. eLife 8 (September). 10.7554/eLife.46750.

Bastian, Amy J. 2006. ‘Learning to Predict the Future: The Cerebellum Adapts Feedforward Movement Control’. Current Opinion in Neurobiology 16 (6): 645–49. 10.1016/j.conb.2006.08.016.

Benwell, Nicola M., Paul Sacco, Geoff R. Hammond, Michelle L. Byrnes, Frank L. Mastaglia, and Gary W. Thickbroom. 2006. ‘Short-Interval Cortical Inhibition and Corticomotor Excitability with Fatiguing Hand Exercise: A Central Adaptation to Fatigue?’ Experimental Brain Research 170 (2): 191–98. 10.1007/s00221-005-0195-7.

Correia José Pedro, João R. Vaz, Christophe Domingos, and Sandro R. Freitas. 2022. ‘From Thinking Fast to Moving Fast: Motor Control of Fast Limb Movements in Healthy Individuals’. Reviews in the Neurosciences 0 (0). 10.1515/revneuro-2021-0171.

Deiber, Marie-Pierre, Manabu Honda, Vicente Ibañez, Norihiro Sadato, and Mark Hallett. 1999. ‘Mesial Motor Areas in Self-Initiated Versus Externally Triggered Movements Examined With fMRI: Effect of Movement Type and Rate’. Journal of Neurophysiology 81 (6): 3065–77. 10.1152/jn.1999.81.6.3065.

Eickhoff, Simon B., Klaas E. Stephan, Hartmut Mohlberg, Christian Grefkes, Gereon R. Fink, Katrin Amunts, and Karl Zilles. 2005. ‘A New SPM Toolbox for Combining Probabilistic Cytoarchitectonic Maps and Functional Imaging Data’. NeuroImage 25 (4): 1325–35. 10.1016/j.neuroimage.2004.12.034.

Emanuel, Aviv, Jasmine Herszage, Haggai Sharon, Nira Liberman, and Nitzan Censor. 2021. ‘Inhibition of the Supplementary Motor Area Affects Distribution of Effort over Time’. Cortex 134 (January): 134–44. 10.1016/j.cortex.2020.10.018.

Friston, K.J., L. Harrison, and W. Penny. 2003. ‘Dynamic Causal Modelling’. NeuroImage 19 (4): 1273–1302. 10.1016/S1053-8119(03)00202-7.

Georgopoulos, Apostolos P, and Adam F Carpenter. 2015. ‘Coding of Movements in the Motor Cortex’. Current Opinion in Neurobiology 33 (August): 34–39. 10.1016/j.conb.2015.01.012.

Georgopoulos, Apostolos P., Andrew B. Schwartz, and Ronald E. Kettner. 1986. ‘Neuronal Population Coding of Movement Direction’. Science 233 (4771): 1416–19. 10.1126/science.3749885.

Hou, Li J., Zheng Song, Zhu J. Pan, Jia L. Cheng, Yong Yu, and Jun Wang. 2016. ‘Decreased Activation of Subcortical Brain Areas in the Motor Fatigue State: An fMRI Study’. Frontiers in Psychology 7 (August). 10.3389/fpsyg.2016.01154.

Hunter, Sandra K., Chris J. McNeil, Jane E. Butler, Simon C. Gandevia, and Janet L. Taylor. 2016. ‘Short-Interval Cortical Inhibition and Intracortical Facilitation during Submaximal Voluntary Contractions Changes with Fatigue’. Experimental Brain Research 234 (9): 2541–51. 10.1007/s00221-016-4658-9.

Jäncke, L., M. Peters, G. Schlaug, S. Posse, H. Steinmetz, and H.-W. Müller-Gärtner. 1998. ‘Differential Magnetic Resonance Signal Change in Human Sensorimotor Cortex to Finger Movements of Different Rate of the Dominant and Subdominant Hand’. Cognitive Brain Research 6 (4): 279–84. 10.1016/S0926-6410(98)00003-2.

Jäncke, L., K. Specht, S. Mirzazade, R. Loose, M. Himmelbach, K. Lutz, and N. Joni Shah. 1998. ‘A Parametric Analysis of the ‘rate Effect’ in the Sensorimotor Cortex: A Functional Magnetic Resonance Imaging Analysis in Human Subjects’. Neuroscience Letters 252 (1): 37–40. 10.1016/S0304-3940(98)00540-0.

Kawashima, R., K. Inoue, M. Sugiura, K. Okada, A. Ogawa, and H. Fukuda. 1999. ‘A Positron Emission Tomography Study of Self-Paced Finger Movements at Different Frequencies’. Neuroscience 92 (1): 107–12. 10.1016/S0306-4522(98)00744-1.

Kluger, Benzi M., Lauren B. Krupp, and Roger M. Enoka. 2013. ‘Fatigue and Fatigability in Neurologic Illnesses: Proposal for a Unified Taxonomy’. Neurology 80 (4): 409–16. 10.1212/WNL.0b013e31827f07be.

Konoike, Naho, and Katsuki Nakamura. 2020. ‘Cerebral Substrates for Controlling Rhythmic Movements’. Brain Sciences 10 (8): 514. 10.3390/brainsci10080514.

Latella, Christopher, Onno Van Der Groen, Cassio V. Ruas, and Janet L. Taylor. 2020. ‘Effect of Fatigue-Related Group III/IV Afferent Firing on Intracortical Inhibition and Facilitation in Hand Muscles’. Journal of Applied Physiology 128 (1): 149–58. 10.1152/japplphysiol.00595.2019.

Liu, Jing Z., Zu Y. Shan, Lu D. Zhang, Vinod Sahgal, Robert W. Brown, and Guang H. Yue. 2003. ‘Human Brain Activation During Sustained and Intermittent Submaximal Fatigue Muscle Contractions: An fMRI Study’. Journal of Neurophysiology 90 (1): 300–312. 10.1152/jn.00821.2002.

Luppino, Giuseppe, Massimo Matelli, Rosolino Camarda, and Giacomo Rizzolatti. 1993. ‘Corticocortical Connections of Area F3 (SMA-Proper) and Area F6 (Pre-SMA) in the Macaque Monkey’. The Journal of Comparative Neurology 338 (1): 114–40. 10.1002/cne.903380109.

Lutz, K., S. Koeneke, T. Wüstenberg, and L. Jäncke. 2004. ‘Asymmetry of Cortical Activation during Maximum and Convenient Tapping Speed’. Neuroscience Letters 373 (1): 61–66. 10.1016/j.neulet.2004.09.058.

Madinabeitia-Mancebo, Elena, Antonio Madrid, Antonio Oliviero, Javier Cudeiro, and Pablo Arias. 2021. ‘Peripheral-Central Interplay for Fatiguing Unresisted Repetitive Movements: A Study Using Muscle Ischaemia and M1 Neuromodulation’. Scientific Reports 11 (1): 2075. 10.1038/s41598-020-80743-x.

Madrid, Antonio, Elena Madinabeitia-Mancebo, Javier Cudeiro, and Pablo Arias. 2018. ‘Effects of a Finger Tapping Fatiguing Task on M1-Intracortical Inhibition and Central Drive to the Muscle’. Scientific Reports 8 (1): 1–10. 10.1038/s41598-018-27691-9.

Madrid, Antonio, Josep Valls-Solé, Antonio Oliviero, Javier Cudeiro, and Pablo Arias. 2016. ‘Differential Responses of Spinal Motoneurons to Fatigue Induced by Short-Lasting Repetitive and Isometric Tasks’. Neuroscience 339 (December): 655–66. 10.1016/j.neuroscience.2016.10.038.

Mahan, Margaret Y., and Apostolos P. Georgopoulos. 2013. ‘Motor Directional Tuning across Brain Areas: Directional Resonance and the Role of Inhibition for Directional Accuracy’. Frontiers in Neural Circuits 7. 10.3389/fncir.2013.00092.

Manjaly, Zina Mary, Neil A. Harrison, Hugo D. Critchley, Cao Tri Do, Gabor Stefanics, Nicole Wenderoth, Andreas Lutterotti, Alfred Müller, and Klaas Enno Stephan. 2019. ‘Pathophysiological and Cognitive Mechanisms of Fatigue in Multiple Sclerosis’. Journal of Neurology, Neurosurgery and Psychiatry 90 (6): 642–51. 10.1136/jnnp-2018-320050.

Marreiros, A.C., S.J. Kiebel, and K.J. Friston. 2008. ‘Dynamic Causal Modelling for fMRI: A Two-State Model’. NeuroImage 39 (1): 269–78. 10.1016/j.neuroimage.2007.08.019.

Maruyama, Atsuo, Kaoru Matsunaga, Nobuyuki Tanaka, and John C. Rothwell. 2006. ‘Muscle Fatigue Decreases Short-Interval Intracortical Inhibition after Exhaustive Intermittent Tasks’. Clinical Neurophysiology 117 (4): 864–70. 10.1016/j.clinph.2005.12.019.

Michely, J., L. J. Volz, M. T. Barbe, F. Hoffstaedter, S. Viswanathan, L. Timmermann, S. B. Eickhoff, G. R. Fink, and C. Grefkes. 2015. ‘Dopaminergic Modulation of Motor Network Dynamics in Parkinson’s Disease’. Brain 138 (3): 664–78. 10.1093/brain/awu381.

Penny, W.D., K.E. Stephan, J. Daunizeau, M.J. Rosa, K.J. Friston, T.M. Schofield, and A.P. Leff. 2010. ‘Comparing Families of Dynamic Causal Models’. Edited by K.P. Kording. PLoS Computational Biology 6 (3): e1000709. 10.1371/journal.pcbi.1000709.

Penny, W.D., K.E. Stephan, A. Mechelli, and K.J. Friston. 2004. ‘Comparing Dynamic Causal Models’. NeuroImage 22 (3): 1157–72. 10.1016/j.neuroimage.2004.03.026.

Pisotta, Iolanda, and Marco Molinari. 2014. ‘Cerebellar Contribution to Feedforward Control of Locomotion’. Frontiers in Human Neuroscience 8 (June). 10.3389/fnhum.2014.00475.

Post, Marijn, Anneke Steens, Remco Renken, Natasha M. Maurits, and Inge Zijdewind. 2009. ‘Voluntary Activation and Cortical Activity during a Sustained Maximal Contraction: An fMRI Study’. Human Brain Mapping 30 (3): 1014–27. 10.1002/hbm.20562.

Rao, S. M., P. A. Bandettini, J. R. Binder, J. A. Bobholz, T. A. Hammeke, E. A. Stein, and J. S. Hyde. 1996. ‘Relationship between Finger Movement Rate and Functional Magnetic Resonance Signal Change in Human Primary Motor Cortex’. Journal of Cerebral Blood Flow & Metabolism 16 (6): 1250–54. 10.1097/00004647-199611000-00020.

Rouiller, Eric M., F. Liang, A. Babalian, V. Moret, and M. Wiesendanger. 1994. ‘Cerebellothalamocortical and Pallidothalamocortical Projections to the Primary and Supplementary Motor Cortical Areas: A Multiple Tracing Study in Macaque Monkeys’. The Journal of Comparative Neurology 345 (2): 185–213. 10.1002/cne.903450204.

Sadato, Norihiro, Vicente Ibañez, Gregory Campbell, Marie-Pierre Deiber, Denis Le Bihan, and Mark Hallett. 1997. ‘Frequency-Dependent Changes of Regional Cerebral Blood Flow during Finger Movements: Functional MRI Compared to PET’. Journal of Cerebral Blood Flow & Metabolism 17 (6): 670–79. 10.1097/00004647-199706000-00008.

Samuel, M. 1997. ‘Evidence for Lateral Premotor and Parietal Overactivity in Parkinson’s Disease during Sequential and Bimanual Movements. A PET Study’. Brain 120 (6): 963–76. 10.1093/brain/120.6.963.

Schlaug, Gottfried, Jerome N. Sane, Venkatesan Thangaraj, David G. Darby, Lutz Jäncke, Robert R. Edelman, and Steven Warach. 1996. ‘Cerebral Activation Covaries with Movement Rate’: NeuroReport 7 (4): 879–83. 10.1097/00001756-199603220-00009.

Scott, Stephen H. 2004. ‘Optimal Feedback Control and the Neural Basis of Volitional Motor Control’. Nature Reviews Neuroscience 5 (7): 532–45. 10.1038/nrn1427.

Sharples, Simon A., Jason A. Gould, Michael S. Vandenberk, and Jayne M. Kalmar. 2016. ‘Cortical Mechanisms of Central Fatigue and Sense of Effort’. PLoS ONE 11 (2). 10.1371/journal.pone.0149026.

Stephan, K.E., W.D. Penny, J. Daunizeau, R.J. Moran, and K.J. Friston. 2009. ‘Bayesian Model Selection for Group Studies’. NeuroImage 46 (4): 1004–17. 10.1016/j.neuroimage.2009.03.025.

Stephan, K.E., W.D. Penny, R.J. Moran, H.E.M. den Ouden, J. Daunizeau, and K.J. Friston. 2010. ‘Ten Simple Rules for Dynamic Causal Modeling’. NeuroImage 49 (4): 3099–3109. 10.1016/j.neuroimage.2009.11.015.

Tanji, Jun. 2001. ‘Sequential Organization of Multiple Movements: Involvement of Cortical Motor Areas’. Annual Review of Neuroscience 24 (1): 631–51. 10.1146/annurev.neuro.24.1.631.

Therrien, Amanda S., and Amy J. Bastian. 2019. ‘The Cerebellum as a Movement Sensor’. Neuroscience Letters 688 (January): 37–40. 10.1016/j.neulet.2018.06.055.

Van Duinen, Hiske, Remco Renken, Natasha Maurits, and Inge Zijdewind. 2007. ‘Effects of Motor Fatigue on Human Brain Activity, an fMRI Study’. NeuroImage 35 (4): 1438–49. 10.1016/j.neuroimage.2007.02.008.

Zénon, Alexandre, Mariam Sidibé, and Etienne Olivier. 2014. ‘Pupil Size Variations Correlate with Physical Effort Perception’. Frontiers in Behavioral Neuroscience 8 (August). 10.3389/fnbeh.2014.00286.

